# BigHeart: Mapping connectivity in the adult human heart at the micron scale

**DOI:** 10.64898/2026.09.11.750845

**Authors:** J. Brunet, A. C. Cook, A. Szmul, M. Use, L. Chestnutt, A. Sivananthan, H. Dejea, T. Urban, A. Bellier, V. Sabarigirivasan, K. Engel, R. Torii, J. Jacob, P. Tafforeau, C. L. Walsh, P. D. Lee

## Abstract

The human heart depends on coordinated muscular, electrical, vascular, lymphatic, and neural systems, but their three-dimensional relationships remain unresolved at microscopic resolution across an intact adult organ. Using hierarchical phase-contrast tomography, we generated a continuous three-dimensional structural reference of a whole adult human heart at 8.01 µm isotropic voxel size, without sectioning or staining. Organ-wide imaging revealed hierarchical myocardial architecture, and quantitative orientation analysis identified regional differences in myocardial cell aggregate organization. The atrioventricular conduction axis was traced into a distinct subendocardial Purkinje network interfacing with working myocardium. Integrated segmentation showed that coronary vessels, lymphatic collectors, and autonomic nerves occupy shared epicardial corridors. This publicly accessible dataset integrates cardiac systems within a common spatial framework for anatomy, computational modeling, and future molecular atlases.

## Main Text

Cardiovascular disease is the leading cause of death worldwide and a major economic burden, costing an estimated €210 billion per year in Europe (*1*) and $320 billion per year in the United States (*2*). Addressing this burden requires a clearer understanding of the structural organization that supports normal cardiac function and is altered in disease. The human heart sustains rhythmic pumping through coordinated electrical activation and mechanical contraction across its four chambers. This coordination depends on the multiscale spatial organization of myocardial cells, the cardiac conduction system, coronary and microvascular networks, valves, nerves and lymphatics throughout the organ. Disruption of this architecture contributes directly to arrhythmias, heart failure and sudden cardiac death (*3*). Yet despite its central importance for function and disease, the complete three-dimensional organization of these systems in the human heart remains incompletely resolved.

Knowledge of cardiac anatomy has been built through classical histology, microdissection, clinical imaging, and regional anatomical studies. These approaches have revealed fundamental features of myocardial cell orientation (*4*), including suggestions of laminar sheet structure (*5*) and the overall trajectory of conduction pathways (*6, 7*). Each method, however, is limited in scope. Histological reconstruction lacks continuity across the full organ (*8, 9*). Clinical imaging cannot resolve cellular or microvascular organization (*10, 11*). Optical and tomographic techniques with near-cellular resolution have thus far been restricted to small animals (*12–14*) or pediatric hearts (*15*). Recent synchrotron imaging has enabled intact adult human organs, including whole hearts, to be imaged in three dimensions (*16, 17*). However, the integrated organization of myocardial, conduction, vascular, lymphatic, and neural systems has not been resolved continuously at microscopic scale within a single intact adult human heart.

In neuroscience, standardized three-dimensional anatomical references (*18–20*) have provided a common framework for linking microstructure to whole-organ function, enabling the field to move from isolated descriptions of local features to integrated circuit-level understanding (*21, 22*). By contrast, the heart lacks a comparable structural reference. Existing cardiac datasets capture macroscale geometry or isolated tissue and cellular systems (*23*–*26*), but not their integrated organization across a single intact adult organ. The absence of such a reference limits the ability to relate microstructural variation to electrical propagation, mechanical contraction and pathological remodeling.

Here, we aimed to create a continuous three-dimensional structural reference of an intact adult human heart that integrates myocardial architecture, the cardiac conduction system, coronary vessels, lymphatics, and autonomic nerves within a common microscopic spatial framework. We used hierarchical phase-contrast tomography (HiP-CT) (*16, 17*), enabled by the Extremely Brilliant Source upgrade of the European Synchrotron Radiation Facility (ESRF) (*27*) to image the complete organ at an isotropic voxel size of 8.01 µm and selected regions at 2.26 µm, without sectioning or staining. Analysis of this reconstruction revealed regional variation in myocardial organization, enabled continuous tracing of the atrioventricular conduction axis, and identified shared epicardial corridors occupied by vascular, lymphatic, and autonomic pathways. The resulting openly accessible reference links chamber-scale anatomy to near-cellular structure and provides a common framework for cardiac anatomy, image-based modeling, and future studies of the heart in health and disease.

### Whole-organ architecture and continuity across scales

We imaged an intact adult human heart from a 63-year-old male donor (donor information in table S1) using HiP-CT at the European Synchrotron Radiation Facility (ESRF) (Fig. 1E; see Materials and Methods). The sample was prepared using a standardized fixation and ethanol-based preservation protocol optimized to maintain structural integrity across scales (*28*). Whole-organ imaging was performed at 8.01 µm isotropic voxel size (Fig. 1E, fig. S1 and Table S2).

**Fig. 1.**
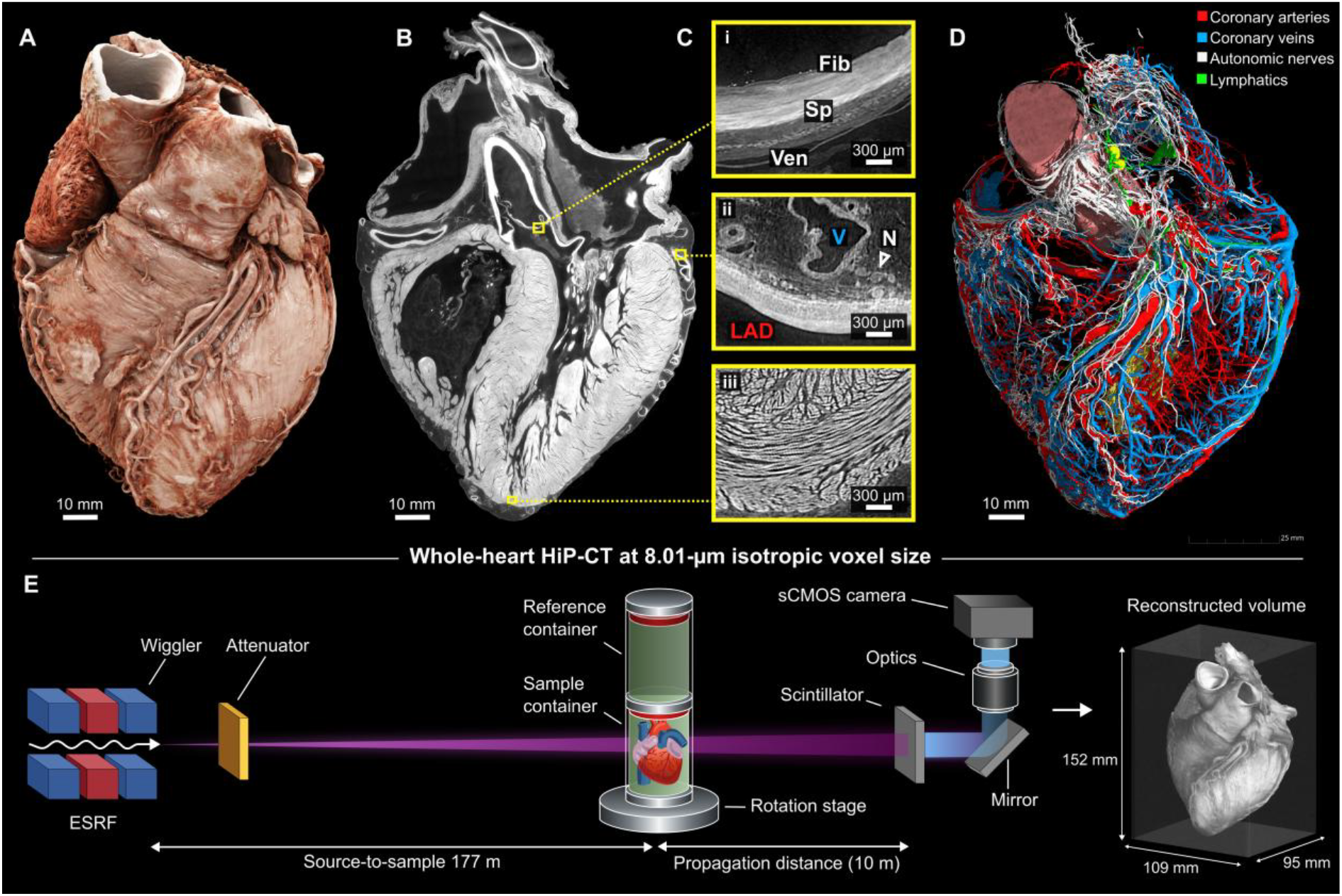
Whole-heart HiP-CT reveals multiscale anatomy in an intact adult human heart. (**A**) Three-dimensional rendering of the intact heart using Cinematic Anatomy (Siemens Healthineers). (**B**) Virtual anatomical section through the 8.01-µm isotropic whole-heart reconstruction, showing chamber morphology, valves, trabeculae, and myocardial structure. Yellow boxes indicate the regions enlarged in (C). (**C**) Representative features resolved within the same volume: (i) Aortic valve leaflet showing the fibrosa (Fib), spongiosa (Sp), and ventricularis (Ven), (ii) an epicardial corridor containing the left anterior descending coronary artery (LAD), an adjacent coronary vein (V), and an autonomic nerve bundle (N, arrowhead), and (iii) apical myocardial aggregate organization. (**D**) Segmented networks, showing coronary arteries (red), coronary veins (blue), autonomic nerves (white), and lymphatics (green) mapped across the heart surface. (**E**) HiP-CT acquisition workflow at the European Synchrotron Radiation Facility (ESRF). The attenuated X-ray beam, generated by a three-pole wiggler, passed through the rotating heart, which was mounted in a sealed sample container beneath a matched reference container on a rotation stage. After 10 m of free-space propagation, the transmitted X-rays were converted to visible light by a scintillator and directed by a mirror through the optics to an sCMOS camera.

After post-processing and cropping, the final dataset had an uncompressed size of 5.7 TB, corresponding to a total volume of 1574 cm^3^ (Fig. 1E). Within this volume, phase-contrast imaging enabled visualization of vascular walls, myocardial aggregates, and smaller branching vessels without sectioning or staining (Fig. 1, A to D). Across the sampled anatomical regions, contrast-to-noise ratios ranged from 3.3 ± 0.5 to 48 ± 8 (table S3).

Segmentation of coronary arteries and veins, lymphatic vessels, and epicardial nerve bundles allowed these anatomically distinct networks to be mapped within the same three-dimensional surface framework (Movie S1, Fig. 1D). Local HiP-CT acquisitions at 2.26 µm isotropic voxel size were also obtained, without sectioning, from selected regions of the intact organ, including the myocardium, conduction axis, and vascular regions (fig. S2 and table S2). These acquisitions resolved finer tissue features and permitted comparison with the corresponding regions in the whole-heart reconstruction. The whole-heart and local datasets are accessible through the Human Organ Atlas (*29*), using Neuroglancer (*30*), an online multiresolution viewer, enabling navigation from the complete organ to local tissue organization (table S4). This continuous spatial framework formed the basis for the subsequent analyses of myocardial architecture, the cardiac conduction system, and epicardial networks.

### Hierarchical organization of myocardial architecture

The ventricular myocardium formed a continuous three-dimensional hierarchy that could be followed from epicardium to endocardium, and from apex to base, in the 8.01 µm whole-heart reconstruction (Fig. 2, A and B). At this scale, the ventricular walls were organized into coherently aligned myocardial aggregates whose transmural orientation changed smoothly and differed between anatomical regions. Local 2.26 µm zoom imaging resolved a finer hierarchy within this architecture, revealing cardiomyocytes joined end to end in branching chains, with these chains grouped into larger sheet-like aggregates that continuously joined and divided.

**Fig. 2.**
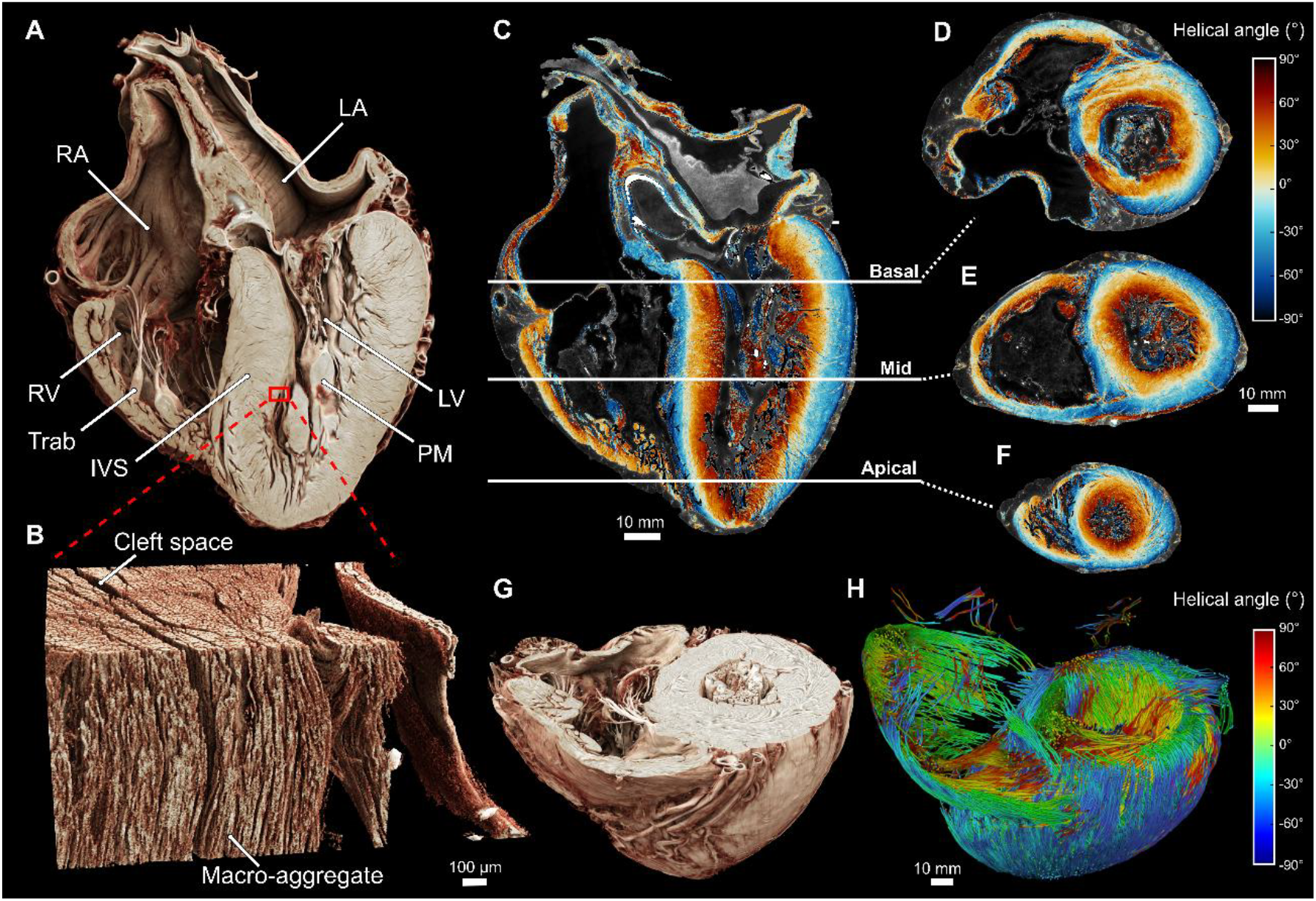
Hierarchical myocardial architecture and aggregate orientation in the intact adult human heart. (**A**), Virtual section through the 8.01 µm isotropic HiP-CT dataset showing the ventricular chambers, septum, trabeculae, and compact myocardium. The red box indicates the region enlarged in (B). (**B**), Ventricular septum showing aligned myocardial macro-aggregates and finer aggregate organization. (**C**), Long-axis section through the heart overlaid with helical angle maps derived from structure tensor analysis, showing transmural variation in cardiomyocyte aggregate orientation from epicardium to endocardium. White lines indicate the basal, mid-ventricular, and apical short-axis levels shown in (D to F). (**D** to **F**), Short-axis helical-angle maps at these levels show transmural rotation and regional differences among the left ventricle, right ventricle, and interventricular septum. (**G**), Three-dimensional rendering of a short-axis myocardial slab showing the ventricular wall, septum, papillary muscles, and trabeculae. (**H**), Tractography of cardiomyocyte aggregate orientation derived from the structure tensor vector field, colored by helical angle, revealing the continuous three-dimensional organization of myocardial architecture across the ventricular walls. IVS, interventricular septum; LA, left atrium; LV, left ventricle; PM, papillary muscle; RA, right atrium; RV, right ventricle; Trab, trabeculae carneae.

Cleft-like spaces contributed to the three-dimensional partitioning of larger aggregates (fig. S2 and Movie S2). These local zooms therefore confirmed, at finer scale, the hierarchical aggregate architecture identified in the corresponding region of the whole-heart reconstruction.

To quantify this architecture across the organ, we applied structure tensor analysis (*31*) to the volumetric data and computed the local orientation of cardiomyocyte aggregates at each myocardial voxel. From these vectors, we derived the predominant angle of cardiomyocyte aggregates, the helical angle (HA), relative to the circumferential axis of the left ventricle (Fig. 2, C - F). As expected, the HA showed a smooth transmural transition, with negative angles in the subepicardium, approaching zero in the mid-wall, and positive values in the subendocardium. An extended 26-segment ventricular model, incorporating nine right ventricular segments alongside the standard 17 left ventricular segments, further illustrated regional variation in myocardial orientation across both ventricles (fig. S3A). The left ventricular free wall and interventricular septum exhibited similarly broad HA distributions, whereas the right ventricular distribution showed a more pronounced peak at positive angles (fig. S3). These findings support the notion that the septum is predominantly left ventricular in architecture. The superior and inferior walls showed differences in fibre orientation, potentially correlating with developmental origins or functional compartmentalization. Notably, the infundibular region showed a layered organization, with an inner trabecular component supporting the pulmonary valve leaflets and an outer compact myocardium. Fractional-anisotropy maps further showed spatial variation in local directional coherence across basal, mid-ventricular, and apical levels (fig. S4). Tractography derived from the same tensor field further emphasized the continuity of aggregate trajectories across the ventricular walls and septum, providing an organ-wide view of the three-dimensional helical arrangement of the myocardium (Fig. 2, G and H). Together, these analyses link local cardiomyocyte and aggregate organization to continuous organ-wide orientation maps, recovering the established left ventricular helical pattern while revealing distinct regional organization in the right ventricle and interventricular septum.

### Three-dimensional morphology of the cardiac conduction system

The atrioventricular (AV) conduction axis could be resolved in continuity from the compact AV node through the non-branching bundle of His, into the left and right bundle branches and into the so-called Purkinje network of the left ventricle (Fig. 3, A to D). The left bundle branch broadened into fascicles that continued distally as a subendocardial Purkinje network. Expert-guided segmentation enabled these specialized pathways to be traced continuously through the surrounding myocardium along the interventricular septum and subendocardial surfaces.

**Fig. 3.**
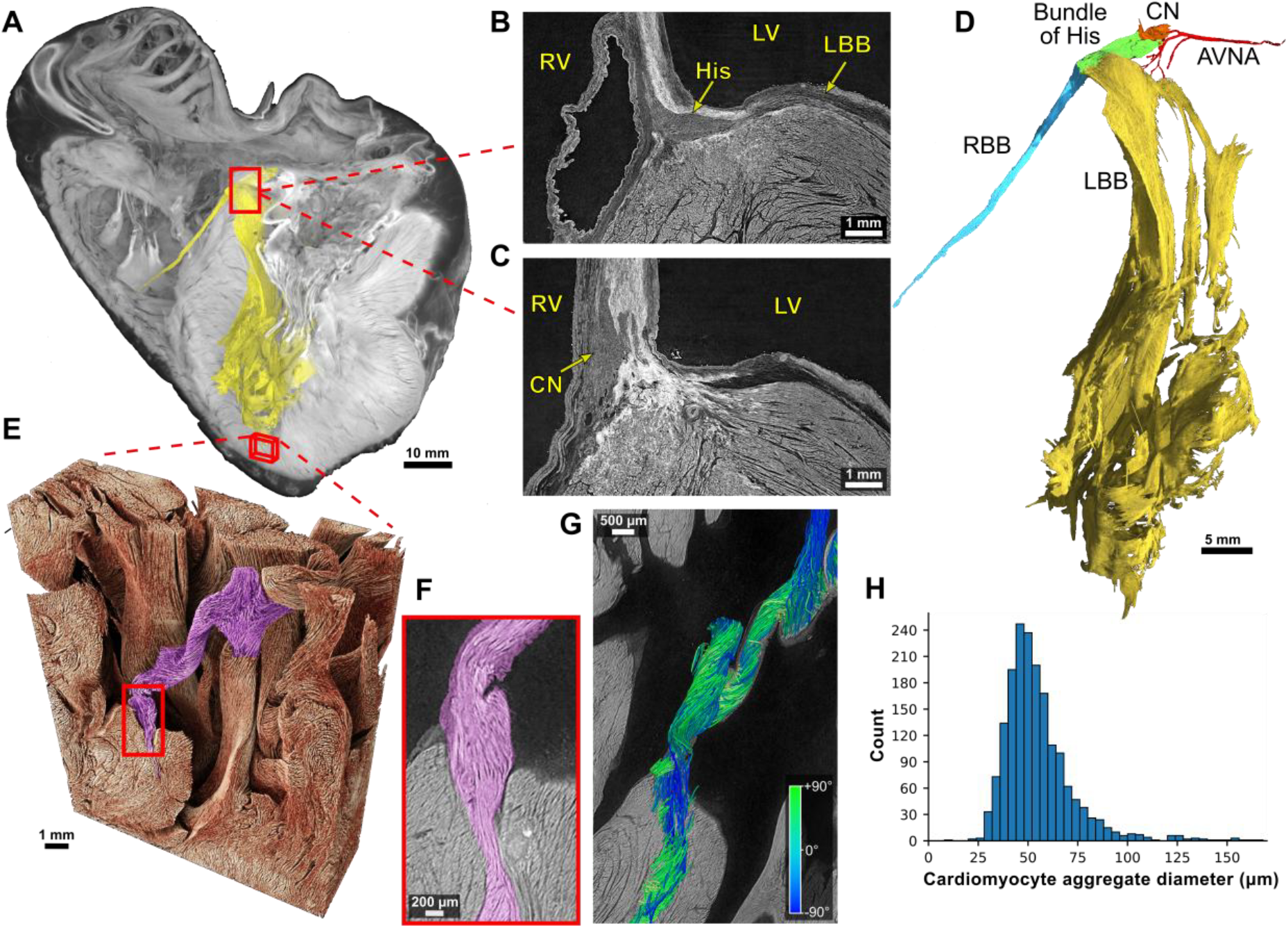
Three-dimensional organization of the cardiac conduction system and a Purkinje-rich trabecula. (**A**), Whole-heart view showing the segmented atrioventricular conduction system in yellow. The analysed trabecula carnea and region of interest are indicated by red boxes. (**B** and **C**), Orthogonal virtual sections through the atrioventricular conduction system. (**D**), Segmentation showing the right (blue) and left (yellow) bundle branches. (**E**), Volume rendering of the trabecula carnea and surrounding myocardium, with the analysed trabecular region highlighted in purple. (**F**), Close-up view of the analyzed Purkinje-rich trabecula, containing predominantly Purkinje fibres together with some working myocardium. (**G**), Cardiomyocyte aggregate centrelines extracted using XFiber extension in Avizo and colored by elevation angle (−90° to +90°). An angle of 0° is parallel to the short-axis plane, and ±90° is perpendicular to it. (**H**) Distribution of cardiomyocyte aggregate diameters measured within the analyzed trabecula carnea. AVNA, atrioventricular nodal artery; CN, compact atrioventricular node; His, non-branching bundle of His; LBB, left bundle branch; LV, left ventricle; RBB, right bundle branch; RV, right ventricle.

Consistent with established descriptions (*6, 32*), the right bundle branch, shown in blue in Fig. 3D, formed a discrete tubular tract that coursed through the septomarginal trabeculation toward the moderator band, whereas the left bundle branch, shown in yellow, exhibited a broad, fan-like morphology. After emerging from the non-branching bundle, it subdivided into anterior and posterior fascicles; the posterior fascicle bifurcated again into two secondary branches that spread across the inferoseptal and inferolateral left ventricular walls. This hierarchical arborization resulted in substantially greater subendocardial coverage in the left ventricle compared to the right.

Distally, the Purkinje network formed a continuous subendocardial mesh that was densest at the left ventricular apex, where fibres penetrated the underlying myocardium at steep angles (Fig. 3, E to H). Its trajectory was not consistently collinear with the local orientation of adjacent myocardial aggregates. Three-dimensional comparison of the local orientation vectors quantified these angular differences across the compact atrioventricular node, His bundle, right bundle branch, basal and mid portions of the left bundle branch, and distal left bundle branch/Purkinje network (fig. S5). These observations indicate that the conduction system forms a distinct, hierarchically branching network embedded within, but not simply aligned with, the surrounding working myocardium. By resolving the conduction pathways together with their surrounding myocardial architecture, this whole-heart dataset provides a three-dimensional anatomical framework for examining chamber-specific activation geometry and the structural relationship between the Purkinje network and working myocardium.

### Epicardial coordination of vascular, lymphatic, and autonomic pathways

Whole-heart segmentations showed that coronary arteries and veins formed the major epicardial scaffold, following the atrioventricular and interventricular grooves before branching across the cardiac surface (Fig. 4A and movie S1). Although fewer lymphatic collectors were identified than coronary vessels or autonomic nerve bundles, the reconstructed lymphatic network showed a clear spatial organization, with collectors frequently coursing alongside major coronary routes, particularly around the cardiac base (Fig. 4, A, B, D, E, and G), consistent with previous descriptions of human cardiac lymphatic anatomy (*33*). Skeletonization of the extracted coronary arterial networks revealed maximum relative Strahler orders of 8 and 7 for the left and right coronary trees, respectively. Vessel diameters and segment counts were quantified by order for each tree (fig. S6). Three-dimensional distance analysis of 6,608 lymphatic collector and 328,074 autonomic nerve centreline points showed that 49.5% of lymphatic points and 60.8% of nerve points were within 2 mm of a coronary arterial surface, increasing to 90.1% and 93.0%, respectively, within 5 mm. Applying the more restrictive criterion of proximity to both coronary arterial and venous surfaces, 40.9% of lymphatic points and 47.5% of nerve points were within 2 mm of both surfaces, increasing to 75.8% and 81.3%, respectively, within 5 mm (fig. S7). These measurements provide organ-wide quantitative evidence that lymphatic and autonomic pathways occupy shared epicardial routes with the coronary vasculature.

**Fig. 4.**
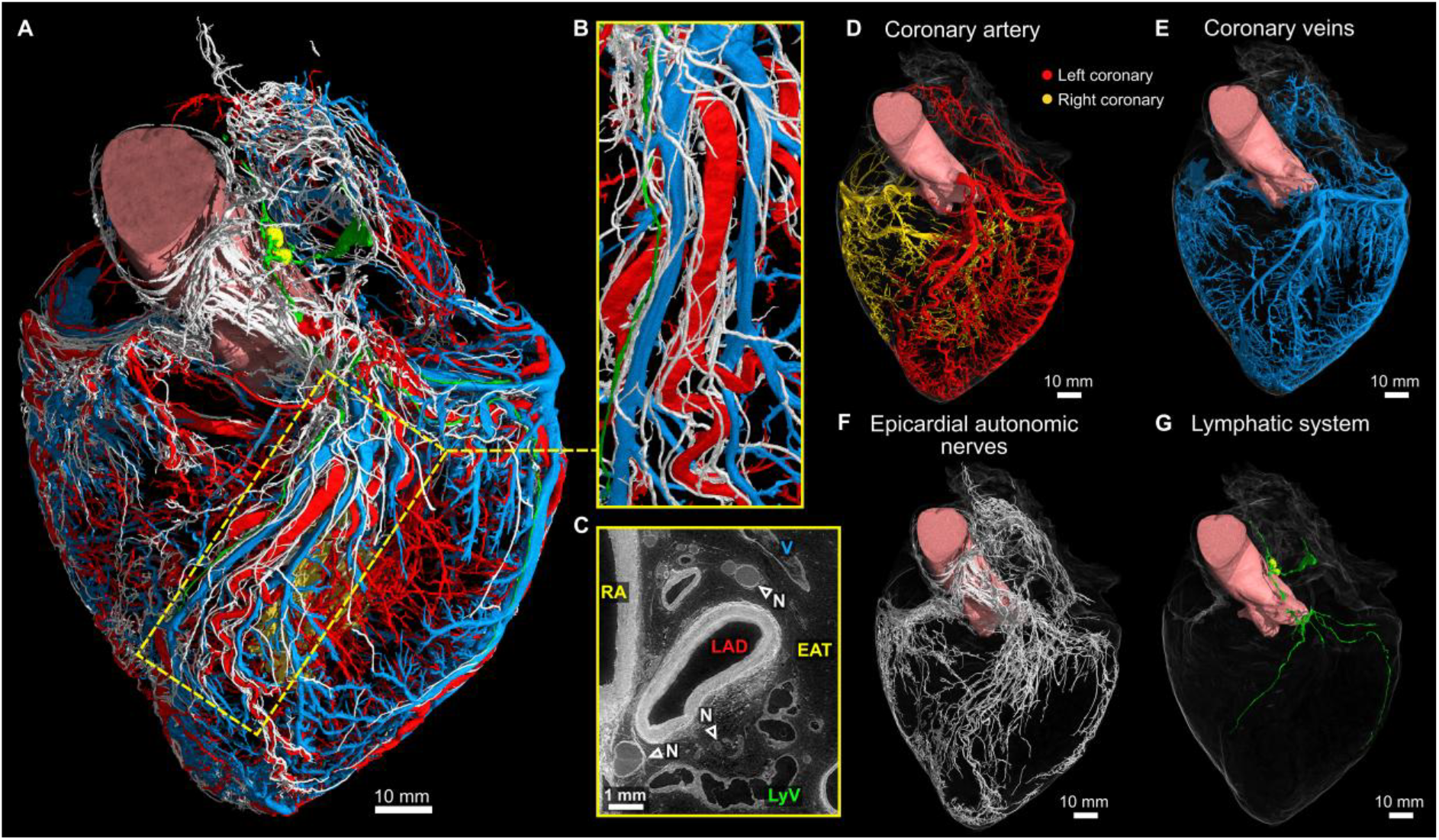
Epicardial coordination of vascular, lymphatic, and autonomic pathways in the intact adult human heart. (**A**), Whole-heart rendering showing the integrated spatial organization of coronary arteries, coronary veins, lymphatic collectors, and epicardial autonomic nerve bundles on the heart surface. Arteries are shown in red and yellow, veins in blue, lymphatic vessels in green, and autonomic nerves in white. The yellow dashed box indicates the anterior interventricular groove region enlarged in (B). (**B**), Local three-dimensional rendering of the same epicardial corridor, showing the close spatial association of coronary vessels, lymphatic collectors, and autonomic nerve bundles. (**C**), Corresponding HiP-CT virtual section through the anterior interventricular groove showing the left anterior descending artery (LAD), right atrium (RA), epicardial adipose tissue (EAT), autonomic nerve bundles (N, arrowheads), and lymphatic vessels (LyV). (**D** to **G**), Separate whole-heart renderings of the coronary arterial tree, coronary venous network, epicardial autonomic nerves, and lymphatic system, respectively, showing that these systems retain distinct branching patterns while occupying shared epicardial routes.

Autonomic nerve bundles were identified within epicardial adipose tissue by their fascicular organization and surrounding epineurium. They were particularly apparent between the aorta and pulmonary trunk and adjacent to the left anterior descending and right coronary arteries, where they coursed alongside coronary vessels within shared epicardial corridors (Fig. 4, A and F), consistent with previous descriptions of human coronary innervation (*34*). Local HiP-CT sections distinguished these bundles from adjacent vessels and adipose tissue in the native grayscale data (Fig. 4C).

Collecting lymphatics could be traced toward two aortopulmonary lymph nodes located between the ascending aorta and pulmonary trunk (Fig. 5). Multiple afferent lymphatic channels entered the nodes, and valves were visible within adjacent collectors. HiP-CT resolved the nodal capsule, cortical regions and follicles, medullary compartments, and internal microvasculature in three dimensions (Fig. 5).

**Fig. 5.**
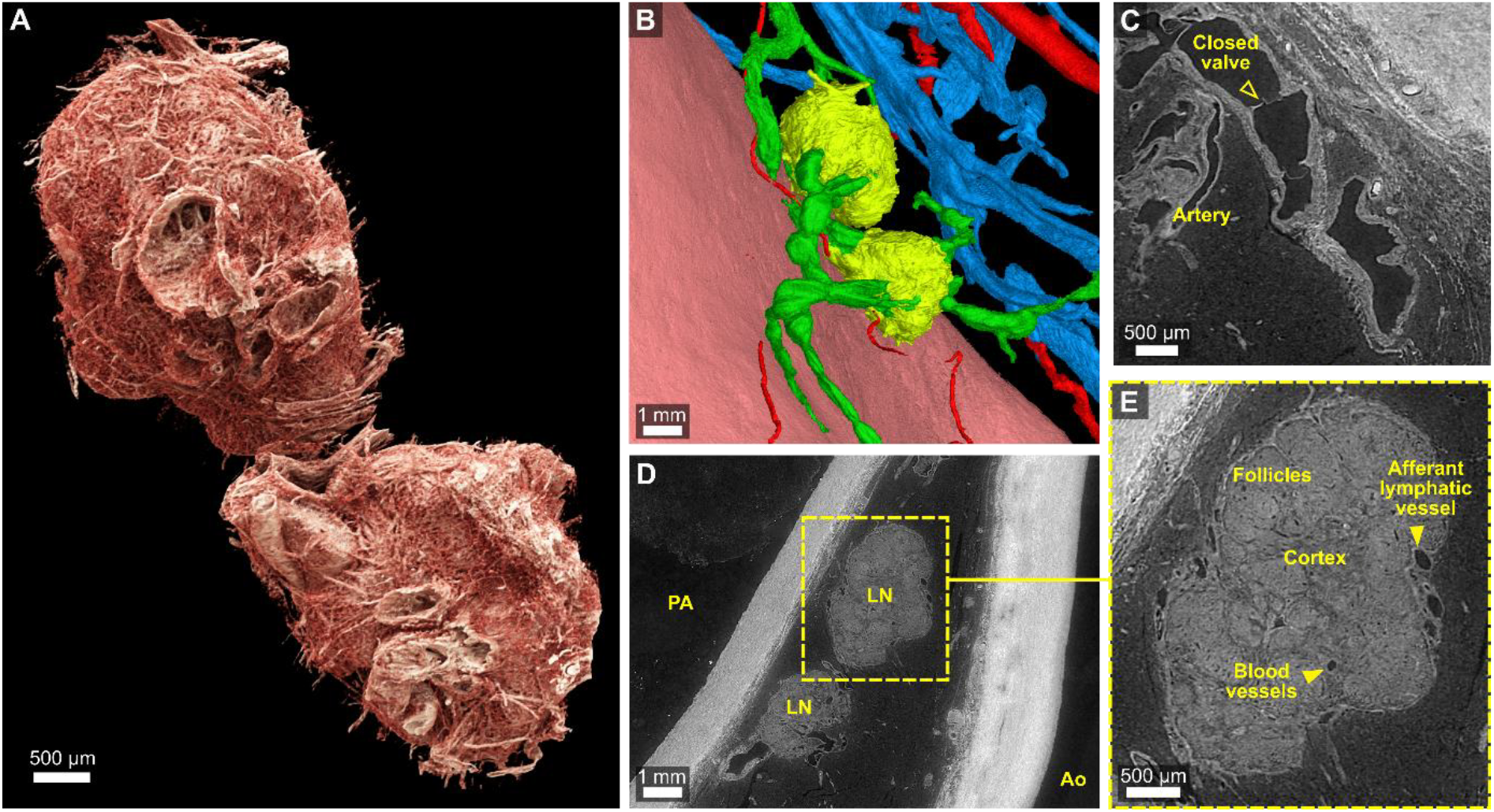
Aortopulmonary lymph nodes and afferent lymphatic drainage resolved in situ by whole-heart HiP-CT. (**A**), Three-dimensional rendering of two aortopulmonary lymph nodes located between the ascending aorta and pulmonary artery. (**B**), Enlarged rendering showing lymph nodes (yellow) connected to collecting lymphatic vessels (green), with coronary arteries (red) and veins (blue) shown for spatial context. (**C**), Virtual HiP-CT section through a collecting lymphatic vessel showing an intraluminal valve adjacent to a coronary artery. (**D**), Virtual section showing the two lymph nodes (LN) flanking the pulmonary artery (PA) near the ascending aorta (Ao). The dashed box indicates the region enlarged in (E). (**E**), Virtual section through a lymph node showing cortical follicles, cortex, internal blood vessels, and an afferent lymphatic vessel, resolved without dissection or contrast agents.

## Discussion

By imaging a complete adult human heart at 8.01 µm isotropic voxel size, with targeted 2.26 µm acquisitions, we established a three-dimensional structural reference integrating myocardial architecture, the atrioventricular conduction axis, coronary vessels, lymphatics, lymph nodes, epicardial nerves, and adipose tissue. Histological and anatomical approaches provide detailed local information (4, 5), whereas clinical imaging and existing cardiac atlases preserve broader anatomical context but lack comparable microscopic detail (10, 11, 23–26). The principal advance of the present work is the ability to examine multiple interconnected cardiac systems continuously within the same intact adult organ.

The myocardial analysis illustrates the value of combining organ-wide continuity with local detail. Unlike diffusion-tensor MRI, which infers orientation from water diffusion, HiP-CT relates the calculated orientation field to visible myocardial aggregate architecture. This relationship enables regional differences in ventricular organization to be interpreted directly alongside the underlying three-dimensional arrangement of myocardial aggregates. The 8.01 µm dataset provides this field across the intact ventricles, whereas the 2.26 µm zoom shows local cardiomyocyte chains joining, dividing, and forming larger aggregates separated by cleft-like spaces (Fig. 2B and fig. S2). These observations agree with regional phase-contrast studies and recent three-dimensional evidence for a hierarchical myocardial mesh (35, 36). They are more readily reconciled with a continuous, heterogeneous mesh than with discrete parallel fibres or a single separable band. This interpretation remains structural, and neither deformation nor force production was measured.

Whole-heart imaging also enabled the atrioventricular conduction axis and distal Purkinje network to be visualized within their surrounding myocardial architecture. The sheet-like morphology of the distal network was consistent with previous descriptions of the human Purkinje system (32), contrasting with the chord-like structures composed of large, rounded and largely sheathed cells originally described by Jan Evangelista Purkinje in sheep. The steep insertion of the distal network into the underlying myocardium and the quantified three-dimensional angular differences between conduction-system and adjacent myocardial orientations (fig. S5) suggest that distal conduction pathways follow trajectories that cannot be inferred solely from the local orientation of working myocardium. These observations support a distal conduction architecture that is morphologically similar to working myocardium yet forms a spatially distinct subendocardial network interfacing with the trabecular myocardium. This structural framework could support future investigations of Purkinje-myocardial connections, ventricular activation models, and the anatomical basis of conduction-system pacing.

Previous anatomical studies described cardiac lymphatic collectors within subepicardial adipose tissue and extensive periarterial autonomic innervation of human coronary arteries (33, 34). Our intact-organ reconstruction extends these observations by placing the coronary vasculature, lymphatic collectors, and autonomic nerves within a common three-dimensional reference and quantifying their spatial proximity across the cardiac surface (fig. S7). Together with the anatomical renderings (Fig. 4), these measurements support the presence of shared epicardial corridors containing vascular, lymphatic, and autonomic structures. The lymphatic collectors could also be traced to aortopulmonary lymph nodes, preserving the three-dimensional continuity of these collecting pathways (Fig. 5).

The public availability of this 5.7-TB dataset is central to its value. The full volume is too large for routine local handling, but browser-based navigation through the Human Organ Atlas website allows exploration across scales without downloading many terabytes of data. This makes the resource useful for validating clinical and diffusion-based imaging, benchmarking segmentation methods, and constraining computational models with measured anatomy.

Several limitations should be considered. The dataset is based on a single adult donor heart. It therefore provides one reference example, but cannot capture anatomical variation across age, sex, body size, or disease. Although no clinically significant cardiovascular disease was recorded, the donor died of pancreatic cancer and had a low body weight; effects of systemic disease, cachexia, or agonal events on tissue dimensions cannot be fully excluded. The heart was also obtained post mortem and subsequently fixed, dehydrated, and imaged ex vivo. These preparation steps were designed to preserve tissue architecture, but some shrinkage or distortion may have occurred, and dehydration may have increased the apparent size of cleft-like spaces between myocardial aggregates. HiP-CT provides structural information, not direct functional measurements of activation, contraction, perfusion, lymph flow, or autonomic control. In addition, identification and segmentation of small structures, particularly distal nerves, lymphatic branches, and Purkinje pathways, remain partly based on morphology and expert anatomical interpretation. Access to synchrotron beamlines capable of imaging intact human organs at this scale remains limited, constraining experimental throughput and the size of cohorts that can be examined, although ongoing improvements in HiP-CT instrumentation and acquisition strategies are progressively reducing imaging times (37). Finally, the size of the raw and reconstructed data remains a practical challenge for widespread quantitative analysis.

Together, these data establish a continuous structural reference in which myocardial, conduction, vascular, lymphatic, and autonomic anatomy can be examined across scales within the same adult human heart. Expanding this approach to additional donors and integrating it with molecular, functional, and clinical measurements will be necessary to move from an individual anatomical reference toward a broader multiscale understanding of the human heart in health and disease.

## Acknowledgments

We sincerely thank the donors and their families for their invaluable contributions. Results from such research can potentially increase humanity’s overall knowledge that can then improve patient care. Therefore, these donors and their families deserve our highest gratitude. We sincerely thank Camille Berruyer, Kathleen Dollman and Vincent Fernandez for assistance with beamline operations. We thank David Stansby and Guillaume Gaisné for their support with data management and uploading, and Samuel Airoldi for assistance with organ extraction and preparation. We further thank Clémence Muzelle, Christophe Jarnias, Filippo Cianciosi, Philippe Carceller, and Alessandro Mirone for their contributions to setup development and improvements. We thank Eugenia Sebastiani-Tofano for providing the heart sketch used in Fig. 1E.

## Funding

The European Synchrotron Radiation Facility for provision of synchrotron radiation facilities at BM18 under proposal number md1290 and md1389.

CZI grant DAF2020-225394 and grant DOI https://doi.org/10.37921/331542rbsqvn from the Chan Zuckerberg Initiative DAF, an advised fund of Silicon Valley Community Foundation (funder DOI 10.13039/100014989)

P.D.L. Royal Academy of Engineering (CiET1819/10).

P.D.L., C.L.W. and JB gratefully acknowledge funding from the MRC (MR/R025673/1).

P.D.L. is a CIFAR-MacMillan Fellow in the Multiscale Human Program.

AC’s research is enabled through the Noé Heart Centre Laboratories which are gratefully supported by the Rachel Charitable Trust via Great Ormond Street Hospital Children’s Charity (GOSH Charity). The Noé Heart Centre Laboratories are based in The Zayed Centre for Research into Rare Disease in Children, which was made possible thanks to Her Highness Sheikha Fatima bint Mubarak, wife of the late Sheikh Zayed bin Sultan Al Nahyan, founding father of the United Arab Emirates, as well as other generous funders.

This research was funded in whole or in part by the Wellcome Trust (209553/Z/17/Z, 227835/Z/23/Z and 310796/Z/24/Z).

JJ was also supported by the NIHR UCLH Biomedical Research Centre and by grant number CZIF2024-009938 from the Chan Zuckerberg Initiative Foundation

## Author contributions

Conceptualization: JB, ACC, PT, CLW, PDL

Methodology: JB, ACC, PT, CLW, PDL

Investigation: JB, ACC, HD, TU, AB, PT, CLW, PDL

Formal analysis: JB, A. Szmul, A. Sivananthan, MU, LC, VS

Visualization: JB, MU, A. Sivananthan, LC, KE

Funding acquisition: PT, CLW, PDL

Supervision: JB, ACC, JJ, PT, CLW, PDL

Writing - original draft: JB, ACC, PDL

Writing - review & editing: All authors

## Data and materials availability

All reconstructed HiP-CT volumes, including the whole-heart dataset and local high-resolution regions of interest, are openly available through the Human Organ Atlas repository (https://human-organ-atlas.esrf.eu). Dataset DOIs are provided in Table S4 and cited in references (*38–46*). Each repository record includes an interactive Neuroglancer viewer, enabling multiresolution exploration directly within a web browser without downloading the complete dataset.

HiP-CT data were reconstructed using custom code written in MATLAB 2017 available at https://github.com/HiPCTProject/Tomo_Recon and the PyHST2 software package (*47*).

The Python package CardioTensor, used for myocyte orientation analysis and tractography, is openly available at https://github.com/JosephBrunet/cardiotensor with documentation at https://josephbrunet.github.io/cardiotensor, and can be cited as Brunet et al. (*31*)

## Supplementary Materials

Materials and Methods

Figs. S1 to S7

Tables S1 to S4

References (47-55)

Movies S1 to S2

