## Supplementary Materials for "BigHeart: Mapping connectivity in the adult human heart at the micron scale"

**The PDF file includes:**

Materials and Methods
Figs. S1 to S7
Tables S1 to S4
References (47-55)

**Other Supplementary Materials for this manuscript include the following:**

Movies S1 to S2

### Materials and Methods

#### Sample preparation

A heart was obtained from a 63-year-old male body donated to the Laboratoire d'Anatomie des Alpes Françaises (LADAF) following the current French legislation for body donation. Available medical information of the organ donor can be found in Supplementary Table S1. Written informed consent was obtained antemortem. All dissections respected the memory of the deceased. Transport and imaging protocols were approved by the Health Research Authority (HRA), the Integrated Research Application System (IRAS) (200429), and the French Health Ministry.

Embalming of the body was performed by injecting formalin into the right carotid artery post-mortem, followed by storage at a temperature of 3.6 °C. During evisceration, the pericardium and lungs were removed. Subsequently, the heart was post-fixed in 4% neutral buffered formalin for a duration of 4 days at ambient temperature. After fixation, the heart underwent a partial dehydration process, transitioning through various ethanol solutions until reaching 70% ethanol concentration. To prevent the bubble formation during imaging, a degassing cycle was applied at each stage using a diaphragm vacuum pump (Vacuubrand, MV2). The heart was then embedded in an agar-ethanol gel for stabilization, oriented with the apex pointing downward. This embedding process included multiple degassing steps to remove any residual air bubbles. The heart was then enclosed in a container and preserved for future scanning. For more information, see Brunet et al. (*28*).

#### Data acquisition

Imaging was performed using hierarchical phase-contrast tomography (HiP-CT) at the European Synchrotron Radiation Facility on the BM18 beamline, which is optimized for propagation-based X-ray imaging of large specimens. Unlike conventional attenuation-based CT, propagation-based phase-contrast imaging exploits differences in the X-ray refractive properties of tissues to enhance soft-tissue contrast. At the ESRF BM18 beamline, the high coherence and energy of the X-ray beam enable these contrast mechanisms to be applied across intact human organs. The heart was imaged using a quasi-parallel polychromatic beam filtered with sapphire and silicon dioxide attenuators, with a source-to-sample distance of 177 m and a propagation distance of 10 m. Whole-organ acquisition was performed at 8.01 µm isotropic voxel size, using 30,000 projections at an average energy of 94 keV, for a total acquisition time of 14.9 hours. X-rays were converted using a 2000 µm LuAG:Ce scintillator (Crytur, Czechia) and detected with an Iris 15 camera (Teledyne Photometrics; 5,056 × 2,960 pixels). To extend the lateral field of view to ~147 mm, a quarter-acquisition scheme was implemented (*17*), combining central and annular scans into a single stitched dataset per stage, as illustrated in Supplementary fig. S1. Each stage covered ~10 mm vertically, and the full heart was acquired using an automated z-series of 25 overlapping stages (6 mm step, ~40% overlap). Flat-field correction was performed using a reference scan acquired from a sealed container filled with 70% ethanol and agar. In addition to the whole-organ dataset, local high-resolution scans at 2.26 µm voxel size were acquired in selected regions of interest, as described previously (*16*).

#### Data processing

The flat-fields, calculated using the reference container scan, were applied to their respective scans. Following stitching of the central and annular scans, the 3D volume was reconstructed using the software PyHST2 (*47*), combining single-distance phase retrieval (*48*), two-dimensional unsharp masking, and filtered back-projection. Subsequently, a modified version of the Lyckegaard algorithm was applied to correct for ring artefacts (*49*). All calculations were carried out on the Networked Interactive Computing Environment (NICE) computing infrastructure at ESRF. The raw imaging data and reconstructed volume occupied 17.2 TB and 12.9 TB, respectively. After cropping, the final reconstructed volume had an uncompressed size of 5.7 TB. JPEG2000 compression was subsequently used to facilitate storage and distribution.

Image-quality assessment

Signal-to-noise ratio (SNR) and contrast-to-noise ratio (CNR) were measured in the postprocessed 16-bit reconstruction using a custom Python tool. Five representative slices were evaluated for each anatomical region. On each slice, circular regions of interest of equal size were manually positioned within the target tissue and within a homogeneous background region. For each slice, SNR and CNR were calculated as:

$$SNR = \frac{\mu_T}{\sigma_B},$$

$$CNR = \frac{|\mu_T - \mu_B|}{\sigma_B},$$

where $\mu_T$ is the mean intensity within the target-tissue ROI, and $\mu_B$and $\sigma_B$ are the mean and standard deviation of intensities within the background ROI, respectively. Results are reported as the mean ± standard deviation across the five slices.

AI Segmentation and rendering

The atrioventricular conduction system, the arterial and venous systems, the lymphatic system, and the nervous system, were initially segmented using VGSTUDIO MAX 3.5.1 software, semi-automatically using a combination of a bounded region growing algorithm and manual segmentation using a Wacom Cintiq QHD27 tablet. The semi-automatic segmentation results of the atrioventricular conduction system, the arterial and venous systems, the lymphatic system, and the nervous system were used to train a set of multiclass AI models using nnU-Net (*50*) as the backbone. The models were trained at 64 µm, 128 µm, and 256 µm resolutions on a high-performance cluster equipped with two NVIDIA A6000 GPUs, and subsequently used to segment data at 64 µm resolution. Predictions from the individual models were combined using majority voting for contested regions. The resulting segmentations were manually reviewed and corrected where needed using VGStudio MAX 3.5.1 and 3D Slicer 5.6.2. The revised segmentations were then resampled and used to retrain the models at 32 µm, 64 µm, and 128 µm resolutions, with inference performed at 32 µm and outputs combined in the same manner as previously described. The corrected results were then used to train a single model at 32 µm, followed by a final round of manual revision. In total, three iterations of active learning were carried out. 3D renderings were performed using Cinematic Anatomy (Siemens Healthineers) (*51*) and VGSTUDIO MAX 3.5.1.

Quantification of cardiomyocytes orientation

Cardiomyocyte aggregate orientation was calculated using Python package CardioTensor (*31*), which implements a 3D structure tensor framework (*52*, *53*). The calculation of the structure tensor involved a two-step process using Gaussian filters. Image gradients were first calculated in the three orthogonal directions using Gaussian derivative filters characterized by the noise scale $\sigma$. The resulting gradient products were then averaged within the local neighborhood using a Gaussian kernel characterized by the integration scale ρ. The noise and integration scales were set to σ = 1 and ρ = 2, corresponding to 8.01 µm and 16.02 µm, respectively. The structure tensor was calculated as:

$$S = K_{\rho}.(\nabla_{\sigma}\mathrm{V}(\nabla_{\sigma}V)^{T})$$

Where $S$ represents the structure tensor, $\nabla_{\sigma}$ denotes the gradient operator with window size $\sigma$, V is the volume of interest, and $K_{\rho}$ indicates the Gaussian kernel with window size $\rho$. Consequently, for each voxel within volume V, the structure tensor S was expressed as:

$$\mathrm{S} = \begin{bmatrix} S_{xx} & S_{xy} & S_{xz} \\ S_{xy} & S_{yy} & S_{yz} \\ S_{xz} & S_{yz} & S_{zz} \end{bmatrix}$$

An eigen-decomposition was then applied to the structure tensor to derive the three eigenvalues $\lambda_1, \lambda_2, \lambda_3$ and their corresponding eigenvectors $\vec{v}_1, \vec{v}_2, \vec{v}_3$. The third eigenvector $\vec{v}_3$ associated with the smallest eigenvalue $\lambda_3$ corresponds to the smallest variation in intensities, which indicates the longitudinal axis of the cardiomyocyte aggregates.

To express orientations in anatomical coordinates, the voxelwise vector field was transformed from Cartesian to a left ventricle (LV) centered cylindrical coordinate system whose long axis was aligned with the LV long axis. Helical angle (HA) was defined as the angle between $\vec{v}_3$ and the local transverse plane, defined by the radial and circumferential directions. The intrusion angle was defined as the angle between $\vec{v}_3$ and the local tangential plane defined by the circumferential and longitudinal directions.

Local coherence of aggregate alignment was quantified using fractional anisotropy (FA), computed from the tensor eigenvalues:

$$FA = \sqrt{\frac{3}{2}}\,\frac{\sqrt{(\lambda_1 - \bar{\lambda})^2 + (\lambda_2 - \bar{\lambda})^2 + (\lambda_3 - \bar{\lambda})^2}}{\sqrt{\lambda_1^2 + \lambda_2^2 + \lambda_3^2}}, \qquad \bar{\lambda} = \frac{(\lambda_1 + \lambda_2 + \lambda_3)}{3}$$

Higher FA indicates stronger local alignment of cardiomyocyte aggregates, while lower FA shows reduced microstructural coherence.

Regional analysis of helical angle

To summarize regional variation in cardiomyocyte aggregate orientation, the ventricular myocardial mask was divided using an extended 26-segment model based on the American Heart

Association 17-segment left ventricular model (*54*). Segments 1 to 17 followed the standard left ventricular convention, whereas nine additional segments, numbered 18 to 26, were used to represent the right ventricular free wall. The interventricular septum, left ventricular free wall, and right ventricular free wall were treated as mutually exclusive regions. For each segment and anatomical region, the axial circular mean HA was calculated using the 180° periodicity of myocardial orientation. Voxelwise HA distributions from −90° to +90° were represented using normalized 256-bin histograms, with adjacent bin values connected by lines for visualization. Histogram counts were normalized to sum to one, and no smoothing or kernel-density estimation was applied. No inferential statistical comparisons were performed because all measurements were obtained from a single heart.

<u>Tractography</u>

Cardiomyocyte bundle tractography was performed using CardioTensor (*31*) from the structure tensor derived $\vec{v}_3$ field. A total of 500,000 seed points were randomly distributed within the myocardium, restricted to regions of high local coherence (FA > 0.4). From each seed, a streamline was propagated bidirectionally (forward and backward) by numerical integration along $\vec{v}_3$ using fourth-order Runge Kutta with trilinear interpolation of the vector field and a step size of 0.5 voxels. Tracking was terminated when FA dropped below 0.15, when the local angular deviation exceeded 60°, or when the streamline exited the myocardial mask. Streamlines were visualized with local HA mapped along each trajectory to illustrate transmural rotation.

<u>Purkinje fibre tracing within trabecula carnea</u>

Fibre trajectories within a trabecula carnea were reconstructed from the HiP-CT volume using the XFiber module in Avizo 3D (Version 2023.2, Thermo Fisher Scientific). The analysed trabecula bundle was selected because it was predominantly composed of conduction tissue consistent with Purkinje fibres and inserted into the compact myocardium at the apical region of the left ventricle. A three-dimensional region of interest encompassing the trabecula and its myocardial insertion was extracted from the 8.01 µm per voxel overview scan. Fibre centrelines were automatically traced using XFiber, with parameters tuned to the signal characteristics and expected dimensions of cardiomyocyte aggregates. Spurious tracks were removed by manual inspection. The resulting fibre network was exported as spatial graphs and visualized by overlaying centrelines on orthogonal slices and volume renderings to illustrate the three-dimensional organisation of the trabecular bundle and its insertion into the myocardium. For visualization, fibres were coloured according to their elevation angle, defined as the angle between the local fibre direction and the short axis plane, enabling quantitative assessment of their 3D orientation within the trabecula and insertion.

<u>Three-dimensional comparison of conduction-system and myocardial orientations</u>

Local orientation vectors were obtained from the three-dimensional structure-tensor field. Before comparison, the vector field was downsampled by a factor of four along each spatial dimension, increasing the voxel spacing from 8.01 to 32.04 µm. Each downsampled voxel therefore represented up to $4^3 = 64$ original voxels. The dominant orientation within each group was calculated treating vectors pointing in opposite directions as the same orientation. This reduced computation time, local noise and spatial oversampling. Vectors with fractional anisotropy below 0.2 were excluded. For each anatomically segmented component of the cardiac conduction system, surface voxels were identified. For each surface voxel, valid surrounding myocardium vectors within a three-dimensional radius of 200 µm were used to calculate the dominant local myocardial orientation. Neighbourhoods containing less than 100 valid myocardial voxels were excluded. The unsigned three-dimensional angular difference between the conduction-system orientation, $v_{CS}$, and the dominant surrounding myocardial orientation, $v_{myo}$, was calculated as:

$$\Delta\theta = cos^{-1}\left(\left|v_{cs}.v_{myo}\right|\right)\frac{180}{\pi}$$

The absolute dot product accounts for the 180° directional ambiguity of structure-tensor orientations. Angular differences therefore ranged from 0°, indicating parallel orientations, to 90°, indicating orthogonal orientations. One angular difference was obtained for each successfully analysed conduction-system surface voxel. Angular-difference distributions were evaluated separately for the compact atrioventricular node, His bundle, right bundle branch, basal and mid portions of the left bundle branch, and the distal left bundle branch/Purkinje network. Distributions were visualized using violin plots and summarized using the median and interquartile range.

Coronary arterial skeletonization and morphometry

The segmented coronary arteries were skeletonized using Avizo 3D (v2023.2) and represented as spatial graphs. The left and right coronary trees were analysed separately, excluding the aorta, with each root node defined at the corresponding coronary ostium. Relative Strahler orders were calculated using custom Python code. Starting from each root node, a breadth-first traversal established graph depth and parent-child relationships. Nodes were then processed from the greatest depth towards the root. Terminal nodes were assigned order 1. Each parent node was assigned the maximum order among its children, increased by one when two or more children shared that maximum order. Each edge inherited the order of its distal node. This ordering describes the hierarchy of the extracted coronary networks, whose terminal segments do not necessarily correspond to anatomical vascular endpoints.

Local vessel radii were estimated from cross-sectional perimeters, assuming that perimeter is less affected by vessel collapse or flattening in the absence of physiological intraluminal pressure. Diameters were calculated as twice the perimeter-derived radius, with one representative diameter obtained per vessel segment. Vessel diameter distributions were summarized using medians and interquartile ranges within each Strahler order. Vessel counts were reported separately for each order, with the left and right coronary trees analysed independently (fig. S6).

Spatial proximity of epicardial networks to coronary vascular surfaces

Binary segmentations of coronary arteries, coronary veins, epicardial lymphatic collectors, and autonomic nerves were analyzed in three dimensions. Lymphatic collector and autonomic nerve masks were reduced to one-voxel-wide centrelines by three-dimensional skeletonization (*55*). Coronary arterial and venous surfaces were extracted from the corresponding binary masks by subtracting a one-voxel binary erosion. For each lymphatic collector or autonomic nerve centreline point, the shortest three-dimensional Euclidean distances to the nearest coronary arterial and venous surfaces were calculated using a k-dimensional tree nearest-neighbour search.

Proximity to coronary arteries was summarized as the fraction of centreline points located within 1, 2, and 5 mm of the nearest arterial surface. Cumulative arterial-distance distributions were generated over distances from 0 to 10 mm (fig. S7E). Spatial localization was visualized using two-dimensional short-axis projections, with centreline points within 2 mm of a coronary arterial surface highlighted (fig. S7, A and B). To assess proximity to both coronary arterial and venous surfaces, the shared-corridor distance was defined as:

$$d_{corridor} = \max(d_{arterial}, d_{venous}).$$

A centreline point satisfied a shared-corridor threshold only when both vascular distances were less than or equal to the indicated threshold. Shared-corridor fractions were calculated at 1, 2, and 5 mm. Joint arterial-venous distance distributions were visualized by binning centreline points according to their distances from the nearest arterial and venous surfaces. Bin values were normalized by the total number of centreline points for the corresponding network and displayed on a logarithmic colour scale (fig. S7, C and D).

The analysis included 6,608 lymphatic collector and 328,074 autonomic nerve centreline points. All distance calculations and corridor classifications were performed using the original three-dimensional coordinates. Results were treated as descriptive measurements from a single heart, and no inferential statistical comparisons were performed.

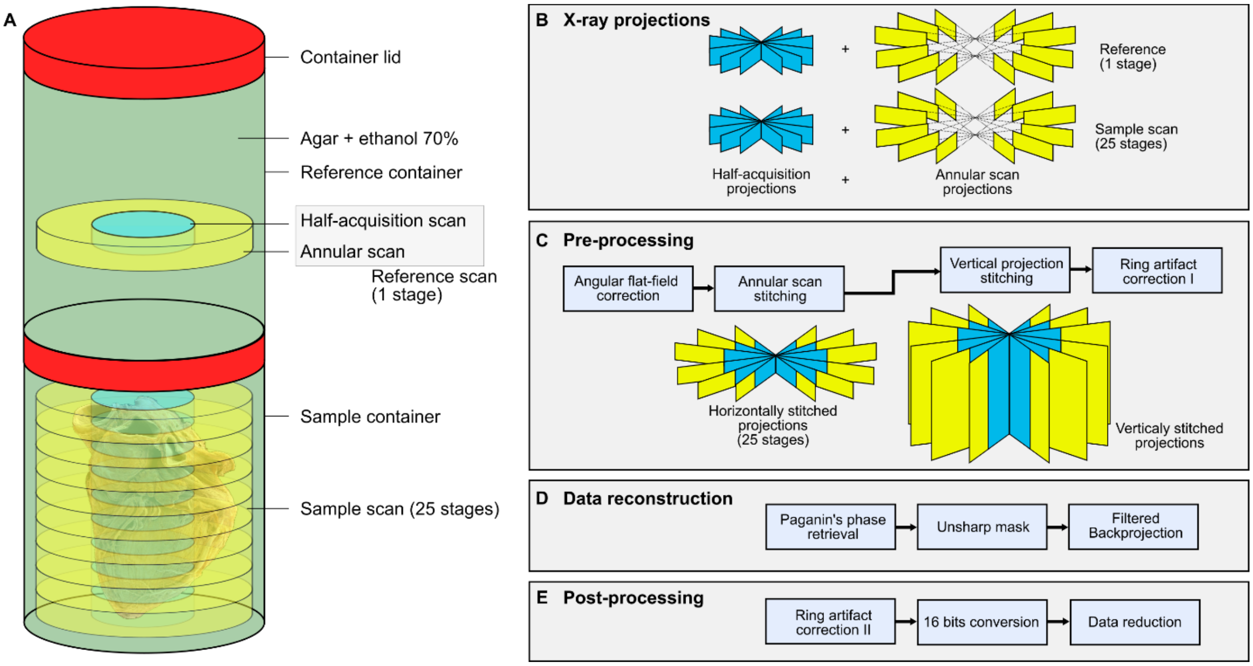


**Fig. S1.**

**HiP-CT scanning strategy and reconstruction pipeline for the 8.01 µm/voxel whole-heart scan.** (**A**) The specimen was immersed in mounting medium in the lower container, while an identical upper container, filled only with the same medium, was used to acquire reference projections for detector and beam normalization. The effective field of view was extended laterally by combining half acquisition and annular scans, and vertically by acquiring a stack of partially overlapping stages along the sample height. (**B**) For each stage, one half-acquisition and one annular scan were acquired. A single reference stage was applied to all sample stages. (**C**) Flat field correction is performed using angular dependent reference projections. Projections from the half acquisition and annular scans are first stitched in the horizontal direction, followed by vertical stitching of consecutive stages. (**D**) Stitched projections underwent single-distance Paganin phase retrieval with an unsharp mask and filtered-backprojection reconstruction. (**E**) Residual ring artifacts are further corrected, intensities are rescaled to a 16-bit dynamic range, and the dataset is reduced for efficient storage and subsequent analysis. Modified from Brunet et al. (16).

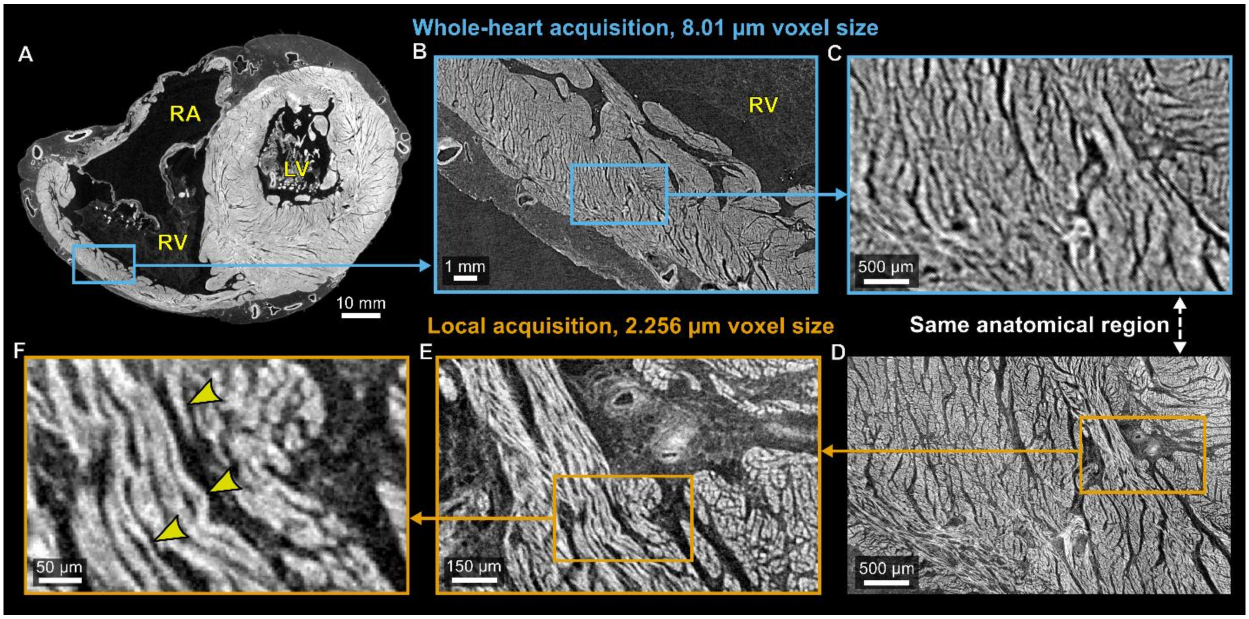


**Fig. S2.**

**Multiscale comparison of myocardial architecture in whole-heart and local HiP-CT acquisitions.** (**A**) Virtual transverse section from the whole-heart dataset acquired at 8.01 µm isotropic voxel size, showing the region selected in the right ventricular myocardium. (**B** and **C**) Successive magnifications of the blue-boxed regions in A and B, respectively. (**D**) The same anatomical field shown in C, acquired separately at 2.256 µm isotropic voxel size. (**E** and **F**) Successive magnifications of the orange-boxed regions in D and E, respectively, revealing finer myocardial aggregate organisation and intervening cleft-like spaces (arrowheads). Yellow arrowheads in F show individual cardiomyocyte bundles. RA, right atrium; RV, right ventricle; LV, left ventricle.

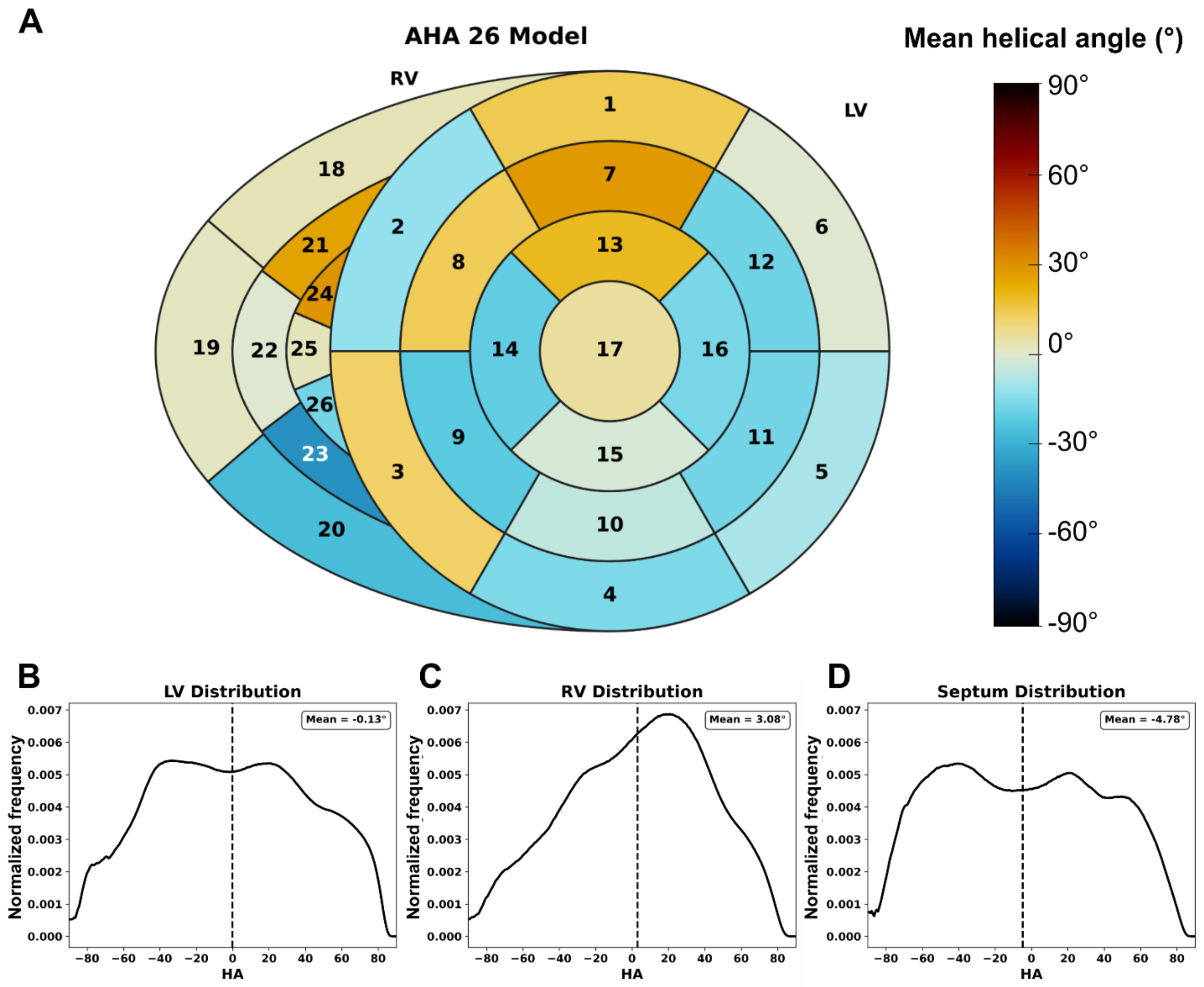


**Fig. S3.**

**Regional helical-angle distributions represented using an extended 26-segment ventricular model.** (**A**) Polar map showing regional mean helical angle (HA) derived from structure-tensor analysis of the 8.01 μm isotropic whole-heart HiP-CT dataset. Segments 1 to 17 correspond to the standard American Heart Association left ventricular model, and segments 18 to 26 extend the model to the right ventricular free wall. Concentric rings represent basal, mid-ventricular, and apical levels, with the central segment representing the apex; individual segments extend through the myocardial wall and do not represent separate transmural layers. Colors indicate the mean voxelwise HA within each segment. (**B** to **D**) Normalized 256-bin histograms of voxelwise HA within the left ventricular free wall, right ventricular free wall, and interventricular septum, respectively. Adjacent bin values are connected by lines for visualization. Vertical dash lines indicate the corresponding mean values. LV, left ventricle; RV, right ventricle.

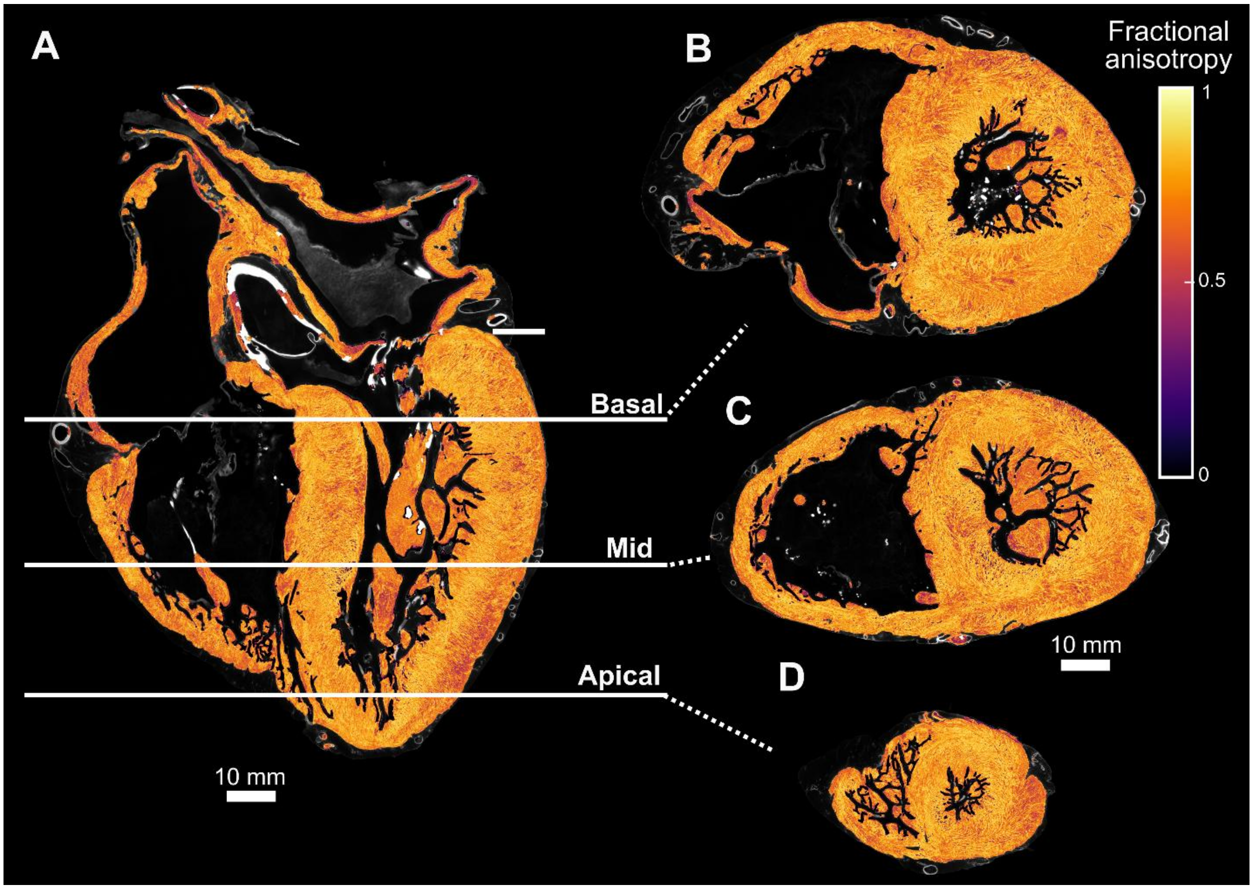


**Fig. S4.**

**Whole-heart distribution of myocardial fractional anisotropy.** (**A**) Long-axis section through the 8.01 µm isotropic HiP-CT dataset overlaid with fractional anisotropy (FA) values derived from structure-tensor analysis. Horizontal lines indicate the basal, mid-ventricular, and apical short-axis levels shown in (B) to (D), respectively. (**B** to **D**) Corresponding short-axis FA maps at the basal, mid-ventricular, and apical levels. FA values range from 0 to 1, with higher values indicating greater local directional anisotropy of the image structure and lower values indicating more isotropic or less directionally coherent structure. Scale bars, 10 mm; the scale bar in (C) applies to (B) to (D).

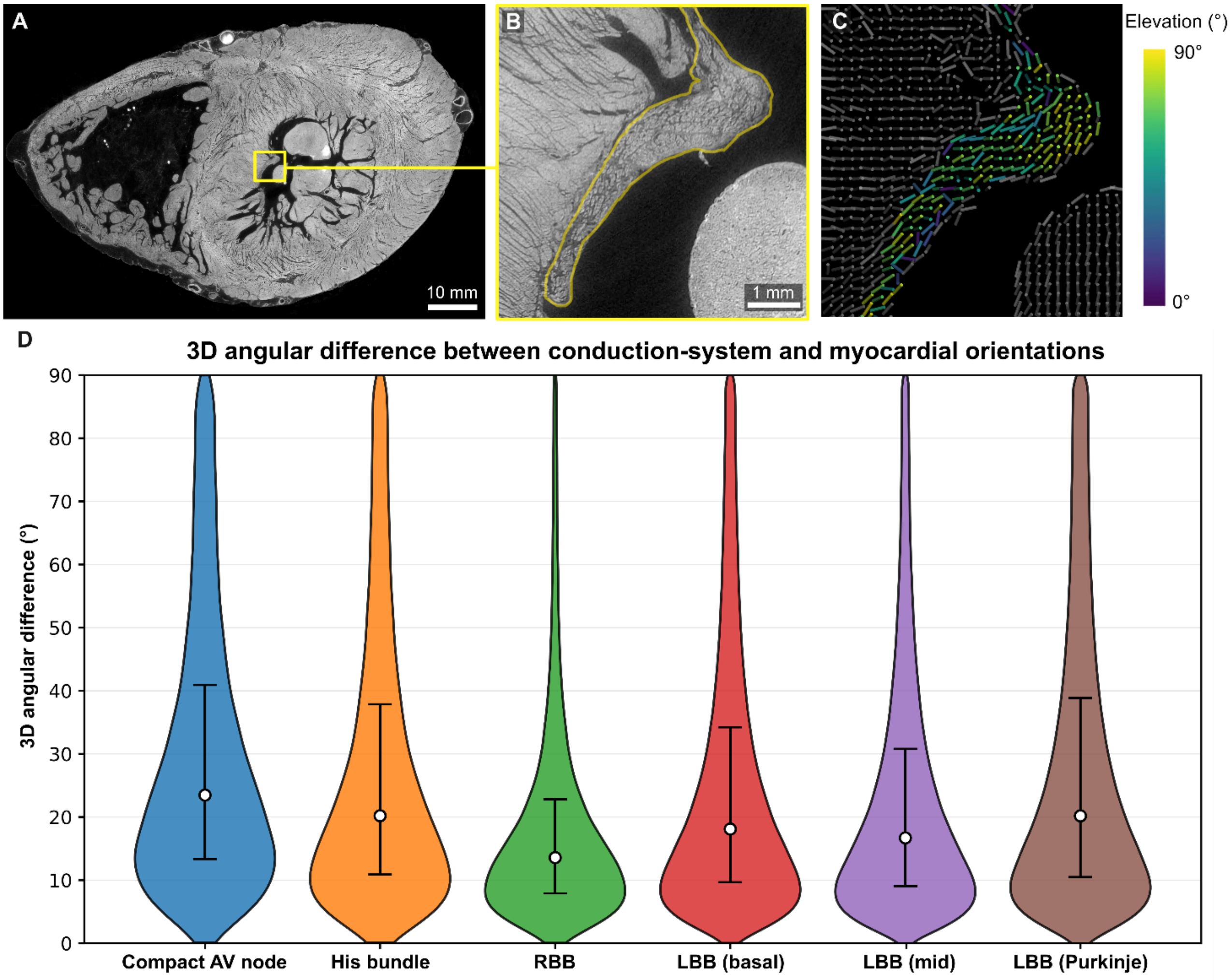


**Fig. S5.**

**Three-dimensional angular differences between the cardiac conduction system and surrounding myocardium.** (**A**) Virtual transverse section through the 8.01-µm isotropic whole-heart HiP-CT dataset. The yellow box indicates the representative region enlarged in (B). (**B**) Magnified virtual section showing a representative portion of the Purkinje network, outlined in yellow. (**C**) Three-dimensional orientation vectors derived from structure-tensor analysis in the region shown in (B). Grey vectors represent adjacent working myocardium, while coloured vectors represent the segmented Purkinje network. Colours indicate the elevation angle of each Purkinje orientation vector relative to the cardiac short-axis plane. (**D**) Violin plots showing the distributions of the unsigned three-dimensional angular difference between conduction-system and local myocardial orientations for the compact atrioventricular node, His bundle, right bundle branch, basal and mid portions of the left bundle branch, and the distal left bundle branch/Purkinje network. A value of 0° indicates parallel orientations, whereas 90° indicates orthogonal orientations. Violin width represents distribution density. White circles indicate medians, and vertical bars indicate interquartile ranges. AV, atrioventricular; RBB, right bundle branch; LBB, left bundle branch.

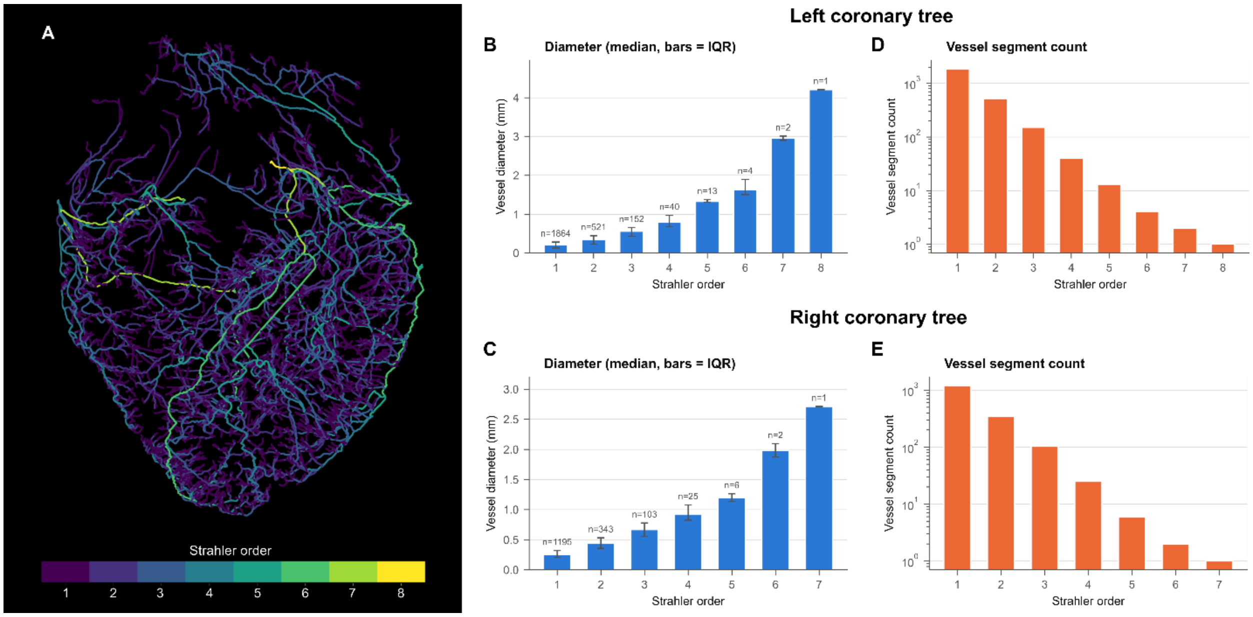


**Fig. S6.**

**Branching hierarchy and vessel diameters of the extracted coronary arterial trees.** (**A**) Three-dimensional coronary arterial skeleton derived from the whole-heart HiP-CT dataset, coloured by relative Strahler order. Extracted terminal segments were assigned order 1, with orders increasing proximally according to the Strahler branching rules. Maximum orders were 8 and 7 for the left and right coronary trees, respectively. (**B** and **C**) Vessel diameter by Strahler order for the left (B) and right (C) coronary trees. Bars indicate medians, error bars indicate interquartile ranges, and *n* denotes the number of vessel segments per order. (**D** and **E**) Corresponding vessel segment counts for the left (D) and right (E) coronary trees, shown on logarithmic vertical axes. Orders describe the hierarchy of the extracted networks, whose terminal segments do not necessarily represent anatomical vascular endpoints.

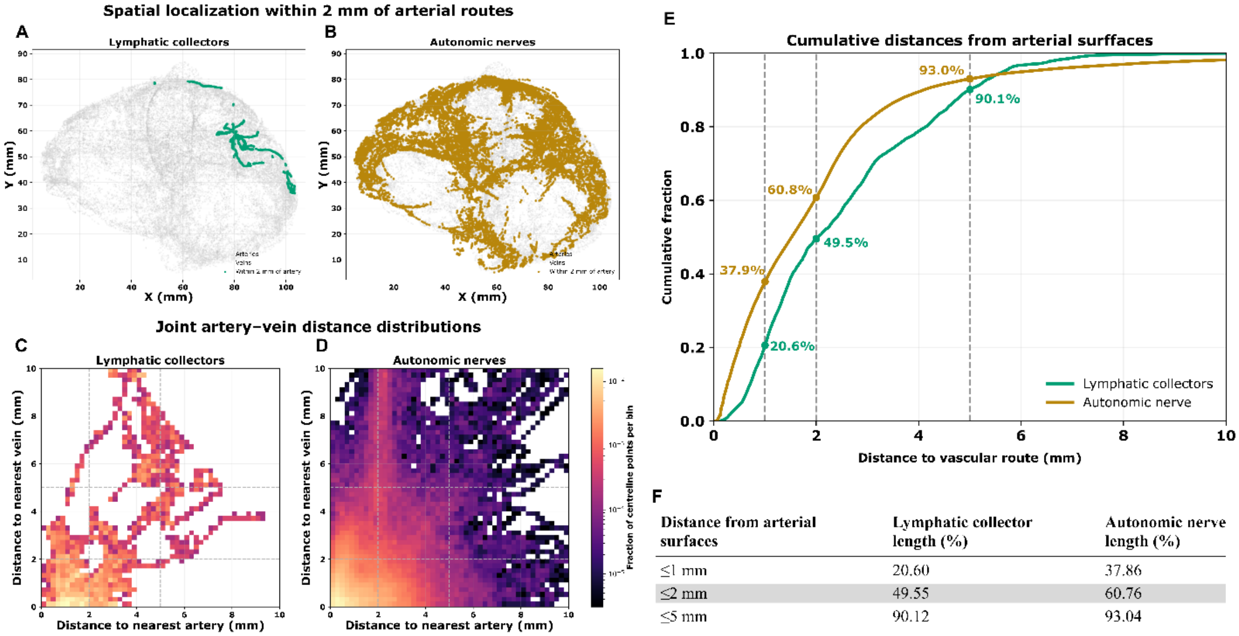


| Distance from arterial surfaces | Lymphatic collector length (%) | Autonomic nerve length (%) |
|---|---|---|
| ≤1 mm | 20.60 | 37.86 |
| ≤2 mm | 49.55 | 60.76 |
| ≤5 mm | 90.12 | 93.04 |

**Fig. S7.**

**Spatial proximity of epicardial lymphatic collectors and autonomic nerves to coronary arterial and venous surfaces.** (**A** and **B**) Two-dimensional short-axis projections of lymphatic collector (A) and autonomic nerve (B) centreline points, in the cardiac short-axis orientation. Coloured points indicate locations within 2 mm of the nearest segmented coronary arterial surface. Arteries, veins, and centreline points outside this criterion are shown in pale grey. All distances were calculated in three dimensions, although two-dimensional projections are shown. (**C** and **D**) Joint distributions of the shortest distances from lymphatic collector (C) and autonomic nerve (D) centreline points to the surfaces of the nearest segmented coronary artery and vein. Colour indicates the fraction of centreline points per bin on a logarithmic scale. Dashed lines indicate the 2-mm and 5-mm thresholds, with the lower-left quadrant corresponding to points within 2 mm of both surfaces. (**E**) Cumulative fractions of lymphatic collector and autonomic nerve centreline points within increasing distance of the nearest segmented coronary arterial surface. Dashed lines indicate distances of 1, 2, and 5 mm, with the corresponding cumulative fractions annotated. (**F**) Percentages of sampled lymphatic collector and autonomic nerve centerline points located within 1, 2, and 5 mm of the nearest segmented arterial surfaces.

**Table S1.**

**Medical information of the body donor.**

| Donor | Age | Sex | Height | Weight | Ethnicity | Year of death | Cause of death | Medical information |
|---|---|---|---|---|---|---|---|---|
| Donor 1, heart | 63 | Male | 178 cm | 60 kg | Caucasian | 2021 | Pancreatic cancer | Additional observations during autopsy: stomach hypertrophy, aspiration of gastric contents. |

**Table S2.**

**HiP-CT acquisition and reconstruction parameters for the whole-heart and local zoom datasets.** Summary of the main acquisition, sample mounting, reference, reconstruction, and post-processing parameters used for the 8.01 µm whole-heart HiP-CT scan and the 2.256 µm local zoom scans.

| Parameter | Whole-heart scan | Local zoom scans (8 zooms) |
|---|---|---|
| **Sample** | LADAF-2021-17 | LADAF-2021-17 |
| **Context** | Whole heart | Volume of interest |
| **Voxel size** | 8.01 µm isotropic | 2.256 µm isotropic |
| **Optic** | DZoom | Zoom |
| **Source** | BM18 lateral beam 1.1 T | BM18 lateral beam 1.1 T |
| **Filters** | sapphire 5.0 mm, $SiO_2$ 30 mm block | Mo 0.23 mm |
| **Average energy** | 94 keV | 104 keV |
| **Propagation distance** | 10.0 m | 2.5 m |
| **Scintillator** | LuAG:Ce 2000 um reflective | LuAG 200 µm |
| **Sensor** | IRIS-15 | PCO edge 4.2 CLHS |
| **Single projection field of view** | 40.5 × 9.6 mm² | 4.62 × 4.62 mm² |
| **Number of projections** | 30000 | 9900 |
| **Exposure time** | 25 ms | 80 ms |
| **Jar mounting** | 140 mm diameter sample container; heart immersed in 70% ethanol and agar | 140 mm diameter sample container; heart immersed in 70% ethanol and agar |
| **Horizontal acquisition scheme** | Quarter acquisition | Half acquisition |
| **Vertical acquisition scheme** | Z-series tomography with 6.0 mm vertical step | Z-series tomography with 3.5 mm vertical step |
| **Reference geometry** | Reference scan acquired from sealed container filled with 70% ethanol and agar | Reference scan acquired from sealed container filled with 70% ethanol and agar |
| **Total field of view diameter** | 147 mm after quarter-acquisition stitching | 8.67 mm |
| **Number of turns** | 25 stages | 6 to 11 stages |

| | | |
|---|---|---|
| **Time per turn** | ~36 min per stage | ~15 min per stage |
| **Total time** | 14.9 h | ~91 to 167 min |
| **Reconstruction protocol** | Flat-field correction using reference scans; central and annular projections stitched; single-distance phase retrieval; unsharp masking; tomographic reconstruction with PyHST2 | Flat-field correction using reference scans; single-distance phase retrieval; unsharp masking; tomographic reconstruction with PyHST2 |
| **Post-processing** | 16-bit conversion, cropping, JPEG2000 compression, and multiresolution binning | 16-bit conversion, cropping, JPEG2000 compression, and multiresolution binning |

**Table S3.**

**Signal-to-noise ratio (SNR) and contrast-to-noise ratio (CNR) measured in selected tissue regions of the 8.01-μm whole-organ HiP-CT reconstruction.** Values are presented as mean ± standard deviation across five slices per region.

| Location | SNR | CNR |
|---|---|---|
| Left ventricle myocardium | 88.87 ± 2.20 | 21.18 ± 0.61 |
| Right ventricle myocardium | 92.33 ± 5.73 | 20.83 ± 1.59 |
| Epicardium | 97.03 ± 6.76 | 3.25 ± 0.45 |
| Papillary muscles | 101.19 ± 5.87 | 26.43 ± 2.65 |
| Aorta | 150.75 ± 25.58 | 47.81 ± 8.44 |

**Table S4.**
**HiP-CT datasets and persistent identifiers.**

| Dataset | Context | Voxel size | Persistent identifier (DOI) |
|---|---|---|---|
| **8.01um_complete-organ** | Whole heart | 8.01 µm | https://doi.org/10.15151/ESRF-DC-2375249432 |
| **2.256um_ROI-1.1** | Local zoom, sinoatrial node anterior | 2.256 µm | https://doi.org/10.15151/ESRF-DC-1659198604 |
| **2.256um_ROI-1.2** | Local zoom, sinoatrial node posterior | 2.256 µm | https://doi.org/10.15151/ESRF-DC-1659198612 |
| **2.256um_ROI-3.1** | Local zoom, left anterior descending coronary artery | 2.256 µm | https://doi.org/10.15151/ESRF-DC-1659198620 |
| **2.256um_ROI-4.1** | Local zoom, atrioventricular node anterior | 2.256 µm | https://doi.org/10.15151/ESRF-DC-1659198628 |
| **2.256um_ROI-4.2** | Local zoom, atrioventricular node middle | 2.256 µm | https://doi.org/10.15151/ESRF-DC-1659198636 |
| **2.256um_ROI-4.3** | Local zoom, atrioventricular node posterior | 2.256 µm | https://doi.org/10.15151/ESRF-DC-1659198644 |
| **2.256um_ROI-5.1** | Local zoom, right ventricular myocardium | 2.256 µm | https://doi.org/10.15151/ESRF-DC-1659302202 |
| **2.256um_ROI-7.1** | Local zoom, interventricular septum | 2.256 µm | https://doi.org/10.15151/ESRF-DC-1659302210 |

**Movie S1.**

**Integrated three-dimensional visualization of the intact adult human heart and its epicardial networks.** Cinematic rendering of the 8.01-µm isotropic whole-heart HiP-CT dataset. Rotation and progressive virtual cropping reveal the external and internal cardiac anatomy and the spatial relationships among the segmented epicardial systems. Coronary arteries are shown in red, coronary veins in blue, lymphatic vessels in green, autonomic nerves in white, and atrioventricular conduction system in yellow. Rendering was performed using Cinematic Anatomy (Siemens Healthineers).

**Movie S2.**

**Three-dimensional visualization of myocardial architecture in the ventricular septum.** Volume rendering of a local region of the ventricular septum from the 2.2 µm isotropic HiP-CT dataset. The movie reveals the hierarchical organization of the human myocardium, from large myocardial macro-aggregates to smaller cardiomyocyte bundles. Cleft-like spaces separate and subdivide these aggregates, exposing a branching three-dimensional architecture that cannot be fully appreciated in two-dimensional histological sections. The near-cellular resolution enables direct visualization of the continuity, alignment, and local reorganization of cardiomyocyte aggregates within the intact adult human heart.
